# Validity of a New Stress Induction Protocol using Speech Improvisation (IMPRO)

**DOI:** 10.1101/2024.09.10.612289

**Authors:** Marina Saskovets, Mykhailo Lohachov, Zilu Liang, Ian Piumarta

## Abstract

This paper proposes a novel stress induction protocol utilizing speech improvisation — Interactive Multitask Performance Response Observation (IMPRO) — in front of an audience. This approach aims to induce stress through a combination of public speaking, cognitive effort, and challenge to maintain a spontaneous narrative. We investigate the validity of this novel approach by examining physiological responses of 35 healthy participants aged 18 to 38. Saliva cortisol was measured as the ground truth for mental stress assessment. In addition, electrodermal activity (EDA) was used as a non-invasive measure of sympathetic activation, offering real-time data with high resolution. Functional near-infrared spectroscopy (fNIRS) measurement was performed to assess the cortical hemodynamic responses induced by the IMPRO protocol. We focused on hemodynamics in the prefrontal cortex (PFC), a region associated with stress processing. We found that cortisol levels and EDA significantly increased in response to the stress task compared to the baseline. We also observed a significant change in PFC hemodynamic levels in a set of channels compared to the baseline, with a higher overall increase in the right frontopolar area compared to the left. In conclusion, our findings validated IMPRO as an effective and easy to perform method for mental stress induction.

## Introduction

Stress is well known for its neuro and biochemical responses. As far as the autonomic nervous system controls other internal organs, stress response involves the whole body through sympathetic activation, increasing cortisol levels, blood glucose, etc. [1]. As Robert Sapolsky noticed, our stress response, initiated by cognitive or social triggers rather than acute survival needs, leads to higher risks of chronic diseases, since it is not adapted for permanent activation [2]. As a result, we can witness a contradictory situation in developed societies where biological stressors such as hunger or physical threats decrease while stress-related abnormalities increase. Goodman et al. notes the need to expand stress research concepts from environmental physiology toward greater concern for perceptual and psychosocial stressors. [3]. Understanding the physiological and psychological effects of stress is crucial for developing effective stress management strategies.

Traditionally, researchers have relied on laboratory methods like the Trier Social Stress Test (TSST) to induce stress in participants, recognized for its effectiveness in creating a stress response through public speaking and social evaluation tasks [4, 5]. While the TSST and its variations, such as the TSST for Groups (TSST-G), are reliable for studying stress, they have limitations in ecological validity as they may not fully replicate real-world stressful situations [6].

Additionally, the TSST does not uniformly induce stress in all individuals due to factors like individual differences in stress resilience or the ability to cognitively disengage from the stressor [7]. Moreover, the structured nature of the TSST, while effective for social stressors,fails to capture the unpredictability and complexity of real-world stressors and can be resource-intensive and logistically challenging to implement, limiting its applicability in large-scale or field studies. Despite these challenges, methods like the Simple Singing Stress Procedure attempt to diversify performance content [8]. The closest to the situation of unpredictability came the developers of the Sing-a-Song Stress Test (SSST) protocol and its variations of anticipatory singing tasks. They provoke stress response by a surprise instruction to sing a song [9, 10]. This is one of the successful and cost-effective options. However, its limitation is its one-time use, whereas the condition of the protocol proposed in this paper assumes a simple algorithm to create reusable unpredictability.

These limitations highlight the need for novel stress induction protocols that are both ecologically valid and efficient to administer. In this study, we proposed a new protocol — Interactive Multitask Performance Response Observation (IMPRO) — that leverages speech improvisation to induce stress. Through improvisation, participants are challenged to think and speak creatively and continuously, replicating the unpredictable and demanding nature of real-life performance. Unlike existing protocols such as SSST, our proposed IMPRO protocol able to create a physiological stress effect regardless of whether the participant is warned. Moreover, improvisation tasks can be easily implemented in a wide range of settings, from laboratory to field, making them a more efficient alternative to some established methods.

In this study, we validate the IMPRO protocol through multimodal measurements, which captures the biochemical, autonomic nervous systematic, and neural responses to the stress induction. Combining multiple streams of physiological signals to study stress has been employed in prior studies [10, 11, 12, 13]. Salivary cortisol levels were measured as the ground truth of the stress level, which is the most common and relevant biochemical indicator of stress widely used in stress studies [14]. Electrodermal (EDA) were measured using a research-grade wearable wristband and serve as indicators of the autonomic nervous system’s response [13, 15]. In addition, cortical hemodynamics were measured using a non-invasive wearable functional near-infrared spectroscopy (fNIRS). Previous research employing the TSST has utilized fNIRS to measure cortical hemodynamic changes during the test, demonstrating its relevance as an indicator of neural responses to stress [16]. By using multimodal measurements, this study intends to provide a comprehensive assessment of the physiological mechanism underlying the stress responses to the proposed IMPRO stress induction protocol.

We would like to pay special attention to the choice of fNIRS as a wearable neuroimaging device. Stress induction in experimental settings has been widely studied using various brain imaging techniques, each offering unique insights into the neural mechanisms underlying stress responses. Functional magnetic resonance imaging (fMRI) has been extensively employed to explore stress-related changes in brain activity, especially subcortical regions such as amygdala, and hippocampus, that allows exploration of the top-down connectivity.

For instance, Veer et al. demonstrated increased functional connectivity between the amygdala and cortical midline structures, specifically the posterior cingulate cortex, precuneus, and medial prefrontal cortex after mental stress [17]. Similarly, electroencephalography (EEG), which offers high temporal resolution, was utilized to examine stress-related changes in brain electrical activity. Recent study on frontal asymmetry, indicated that exposure to stressful environment such as workplace noise led to decreased alpha rhythms, reflecting increased activity in the prefrontal cortex, as well as significant right frontal activation [18]. Despite the valuable contributions of fMRI and EEG, each method has its limitations. FMRI, while providing high spatial resolution, is often constrained by its low temporal resolution and the requirement for participants to remain motionless within a scanner, which may not fully capture the dynamics of stress responses in more naturalistic settings. EEG, on the other hand, offers excellent temporal resolution but lacks the spatial precision needed to localize neural activity with accuracy and shows vulnerability to motion artifacts [19].

Functional near-infrared spectroscopy emerges as a promising alternative that addresses some of these limitations. fNIRS provides a balance between spatial and temporal resolution, making it well-suited for studying cortical hemodynamics in response to stress [20]. Unlike fMRI, fNIRS allows for greater flexibility and mobility, enabling the assessment of brain activity in more ecologically valid environments [21]. Additionally, while fNIRS shares some of the temporal advantages of EEG, it surpasses EEG in its ability to measure localized cortical oxygenation and blood flow changes, offering more specific insights into the regional hemodynamic responses to stress [22]. Recent studies utilizing fNIRS, such as those by Molina-Rodríguez et al., have demonstrated its efficacy in capturing prefrontal cortex activity during social evaluative threats, underscoring its utility in mental stress research [23].

While traditional methods like fMRI and EEG have significantly advanced our understanding of the neural correlates of stress, wearable fNIRS presents distinct advantages in studying stress in more naturalistic and dynamic contexts. Its ability to provide localized measurements of cortical hemodynamics, combined with its flexibility in experimental design, makes it a valuable tool in our research.

## Materials and methods

### Ethics statement

The Kyoto University of Advanced Science’s “Ethical Review for Research on Human Subjects Committee” has approved the study. Approval number 23E03, date of approval 2023/07/13. All participants signed a written informed consent form before taking part in the experiment. Recruitment of participants for this study took place from 13 June 2023 to 15 February 2024, including advertising, pre-screening and experiments.

### Participants

For this study, we included healthy English-speaking adults aged 18 to 35 fluent in English, with normal hearing and no history of neurological or psychiatric disorders. Exclusion criteria were chronic illnesses, current medication affecting cortisol levels or cognitive function, substance abuse, and recent major life stressors. Participants were recruited through a combination of methods to ensure a diverse and representative sample. We posted advertisements and distributed flyers on campus. Additionally, we used word of mouth by informing colleagues and encouraging them to refer potential participants. Out of 44 applicants, 37 participants met the criteria and were enrolled in the study. Seven participants dropped out due to health issues (1), ineligibility discovered during preliminary assessments (2), failure to keep appointments (4). Two more participants were enrolled in the study, but their results were excluded from statistical analysis, due to poor signal quality, or due to session interruption for technical reasons. Participants received a reward (Amazon gift card) to thank them for their efforts and time. Demographic information of participants was assessed and organized into Table 1, including distribution on age, gender, dominant hand, Munich Chronotype Questionnaire (MCTQ), and perceived stress scale (PSS).

**Table 1.** Demographic profile table.

| Demographic | Description | Number | Percentage |
| --- | --- | --- | --- |
| **Age** | 18-23 | 21 | 60,00% |
|  | 24-29 | 8 | 22,86% |
|  | 30-38 | 6 | 17,14% |
|  | Total | 35 | 100,00% |
| **Gender** | Male | 19 | 54,29% |
|  | Female | 16 | 45,71% |
|  | Total | 35 | 100,00% |
| **Dominant hand** | Right | 32 | 91,43% |
|  | Left | 1 | 2,86% |
|  | Ambidexter | 2 | 5,71% |
|  | Total | 35 | 100,00% |
| **Chronotype (MCTQ)** | Morning | 3 | 8,57% |
|  | Evening | 24 | 68,57% |
|  | Night | 8 | 22,86% |
|  | Total | 35 | 100,00% |
| **Perceived stress scale (PSS)** | Low | 9 | 25,71% |
|  | Moderate | 20 | 57,14% |
|  | High | 6 | 17,14% |
|  | Total | 35 | 100,00% |

### Study Procedure

This study uses a within-subject laboratory-based design to evaluate the IMPRO stress induction protocol utilizing multimodal physiological measurements (cortisol, EDA, fNIRS). As shown in Fig. 1, the experiment procedure consisted of six stages: preparation, awake resting or baseline (O-S), anticipation (S-A), IMPRO stress induction (A-D), stress recovery (D-E), and debriefing. In turn, IMPRO stress induction includes three tasks: free improvisation (A-B), random words challenge (B-C), and arithmetic load (C-D). Salivary cortisol levels were used to assess the average efficiency of the IMPRO tasks compared to the baseline. Patterns of EDA and PFC activity were measured and analyzed to assess the average efficiency of the IMPRO tasks compared to baseline and recovery, and dynamic changes between stages.

**Fig 1.**
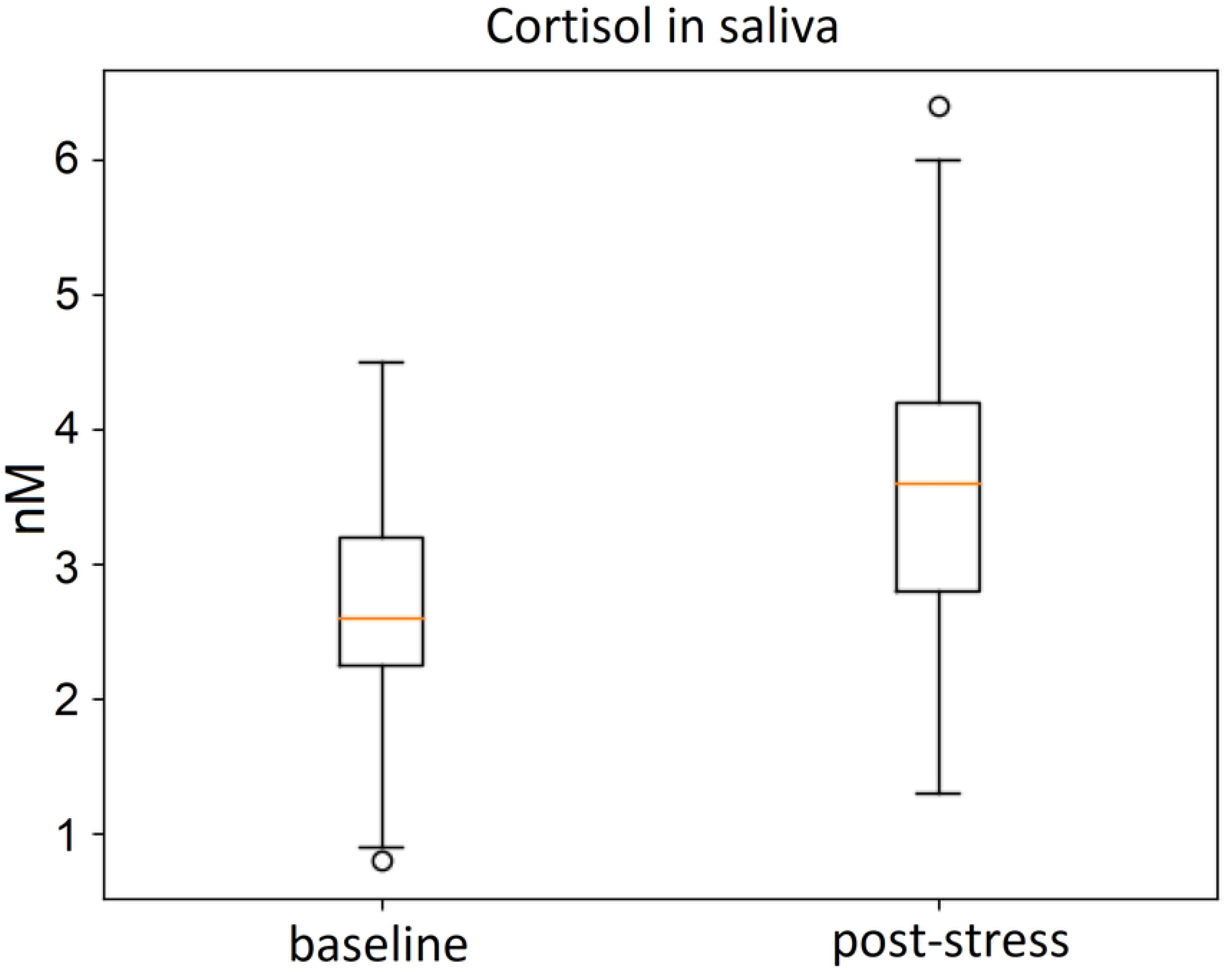
Study Procedure.

#### Setup

The experiment commenced with a preparation stage lasting 15 minutes, during which the researcher welcomed the participant, provided an overview of the study, and ensured they were comfortable. The participant was then asked to sign an informed consent form and to put on all necessary measurement devices, including the wearable wristband and the fNIRS device. They had been informed in advance about the tasks they would undertake.

Following this, the awake resting stage began, where the participant sat quietly for 5 minutes to establish baseline brain and body signals. Next, in the anticipation stage, which also lasted 5 minutes, the participant received detailed instructions about the upcoming improvisation tasks and had the opportunity to ask clarifying questions or prepare their thoughts in silence.

#### Stress induction

Stress induction was provided by improvisation tasks which lasted 10 minutes. This stage consisted of three tasks: free improvisation, random words challenge, and an arithmetic load. Two listeners were present throughout these tasks, and video recordings were made for further analysis. Proposed protocol was presented to participants as a game. The game consists of three tasks with increasing levels of difficulty.

Task A requires the participant to talk without pause on any free topic and lasts 3 minutes. If the participant makes a pause between words for more than 5 seconds, the game starts from the beginning. The purpose of this stage is to give the participant a chance to settle into their performance strategy and warm up.

Task B keeps improvising with the additional task and lasts 4 minutes. In this stage, listeners “throws in” (pronouncing aloud) random words and the speaker incorporates these words into the narrative without pausing to think. The challenge is verbal confusion. This is a divergent type of task, with no right or wrong answer. Multiple strategies for interaction with the new information are available. Participants do not know these words in advance and have to experiment with their strategy on the way. Researcher makes notes regarding the participant’s strategy for interacting with the words (formal repetition, inclusion in the narrative, or ignoring). Even if the participant ignores the words, this is not a reason to stop the game as long as the participant continues to speak.

Task C requires the participant to speak without pauses with the additional task and lasts 3 minutes. The audience “throws in” arithmetic tasks and the speaker’s task is to give the answer and continue the speech immediately, without pausing to think. This is the arithmetic confounding factor of the convergent type (there is one correct answer). Answering incorrectly or ignoring a problem is not a reason to stop the game. The audience does not comment on the accuracy of the answers. An additional stress factor during all 3 tasks is that listeners are not allowed to show signs of non-verbal support and approval (smiling or nodding).

#### Recovery

After completing all three tasks, the participant relaxed in a comfortable chair for 5 minutes. In the recovery stage, participants were encouraged to sit in a more comfortable position while having an informal conversation with the researcher in chief. They involved in a short semi-structured interview in a casual tone about their subjective experience during the tasks: “How was it for you?”, “What did you feel?”, “What was difficult?”, “What was noticeable?”. This stage was used as a second control period to compare an improvisation talk to a casual conversation.

Finally, in the 10-minute debriefing phase, the researcher removed the measurement devices, provided feedback to the participant, and addressed any questions about the study, while also seeking feedback on their experience with the experiment design, the comfort and functionality of the measurement devices, their interactions with the listeners, and their overall feelings about the study.

### Measurements

To validate the stress response during the new IMPRO stress induction protocol, we analyzed changes in brain and body activity in several modalities. We compared cortisol levels — the gold standard biochemical marker of stress — before and after the stress task. We also measured the level of electrodermal activity associated with the autonomic nervous system response. The brain response was examined by measuring hemodynamic changes in the prefrontal cortex. In what follows we describe the measurement systems and devices.

#### Salivary cortisol kit

The SOMA CUBE system was used to measure salivary cortisol levels. SOMA test is based on the principle of lateral flow, also called immunochromatographic strip (ICS) tests. The test was carried out by adding the sample (buffer/saliva mixture) onto the sample pad. As the mixture flowed across the membrane, the cortisol was captured by the test line, resulting in the appearance of a red line. The SOMA Reader measured the line intensity and converted these values into the corresponding cortisol concentrations in the saliva sample in the unit of nM.

In our experiment, saliva samples were collected at two points during the experiment: (1) baseline, after the awake resting phase and before the experimental intervention, and (2) stress, after the IMPRO stress task. To avoid circadian rhythm interference, saliva samples were taken approximately at the same time (between 2 pm and 4 pm) in all experiment sessions. Cortisol levels at each predefined time point were obtained using the SOMA CUBE LFD Reader according to the manual. The data were logged in an Excel spreadsheet.

#### Wearable wristband

The Empatica E4 wristband, an FDA-approved research-grade wristband, was used to measure EDA. Our study focused on the general tonic-level EDA, which relates to the slow activity reflecting general changes in autonomic arousal. Measurements were conducted by noninvasive contact with the skin on both wrists. Participants pressed the time stamp buttons on the bracelet following the researcher’s instructions. Data were measured in the unit of microsiemens (μS), with a sample rate of 4 Hz.

#### Functional Near-Infrared Spectroscopy (f-NIRS)

Cortical hemodynamics were measured using a wearable Aritinis Brite 24 system with a sampling rate of 50 Hz. Optodes were arranged following the 2×12-channel template. The optodes were fixed on a soft neoprene head cap and were distributed symmetrically on the right and left sides of the frontal area. Half of the optodes were placed between the FpZ-F7-F3-FZ region, and another half - between the FpZ-F8-F4-FZ region according to the international 10–20 EEG system (Fig 2, 3). We focus on fNIRS sensor recorded activity in regions of the PFC, which approximately correspond to Brodmann areas 9, 10, and 45 (or the anterior and dorsolateral PFC). This configuration makes it possible to compare activity in the two hemispheres and tracking frontal asymmetry as an indicator of mental stress [24, 25]. The Brite 24 device was connected to the OxySoft via Bluetooth and data were constantly streamed to the OxySoft. The fNIRS system allows the participants to move freely and thus ensure the ecological validity of the measurement. Raw data from wearable devices were collected using complementary software and further imported for preprocessing and statistical analysis, for which Python 3.10 was used.

**Fig 2.**
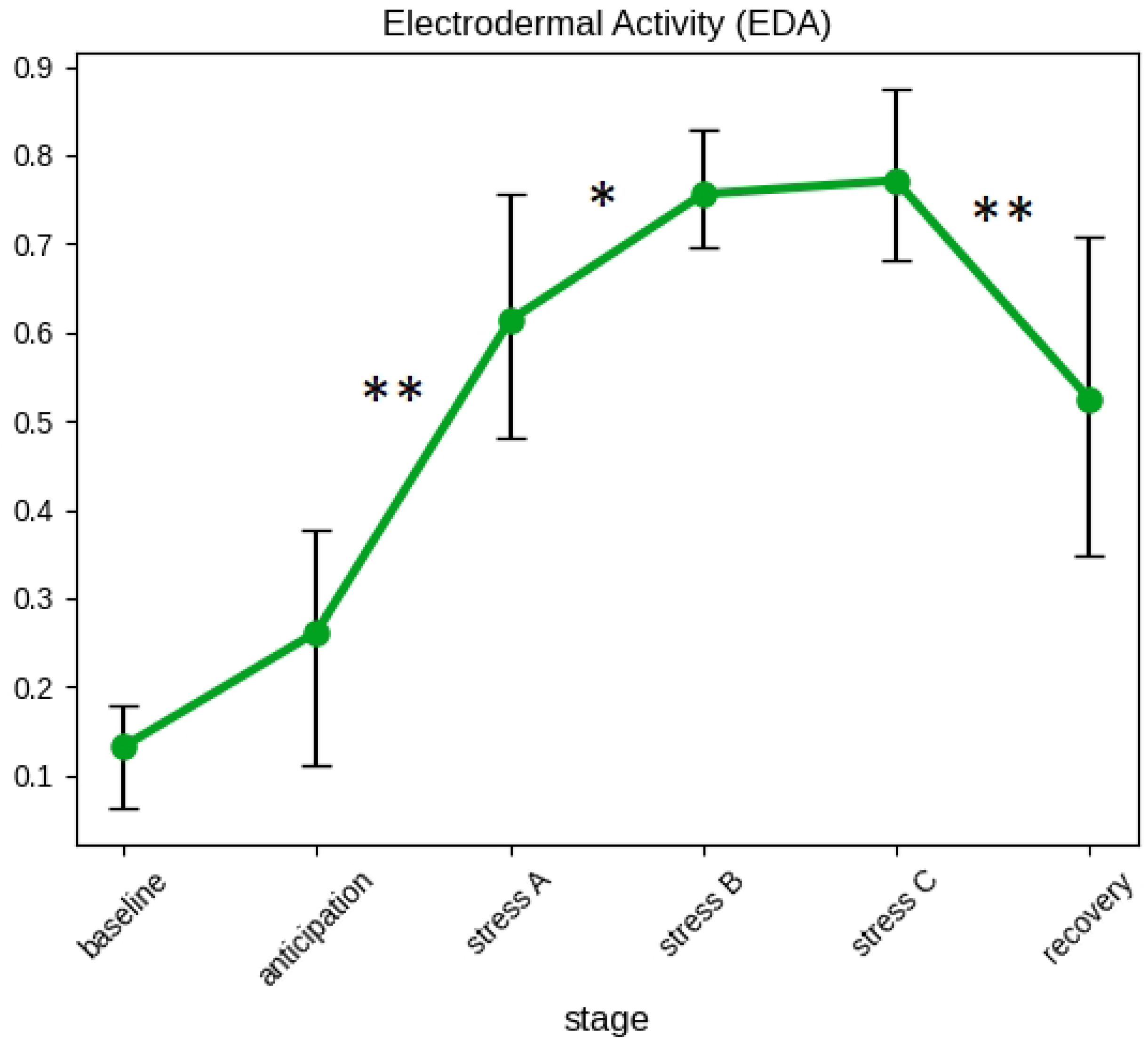
Location of the fNIRS transmitters, receivers, and channels. a) fNIRS device position on the head. b) Channel’s location. The numbered black dots represent the channels in between the transmitters (yellow dots) and receivers (violet dots).

**Fig 3.**
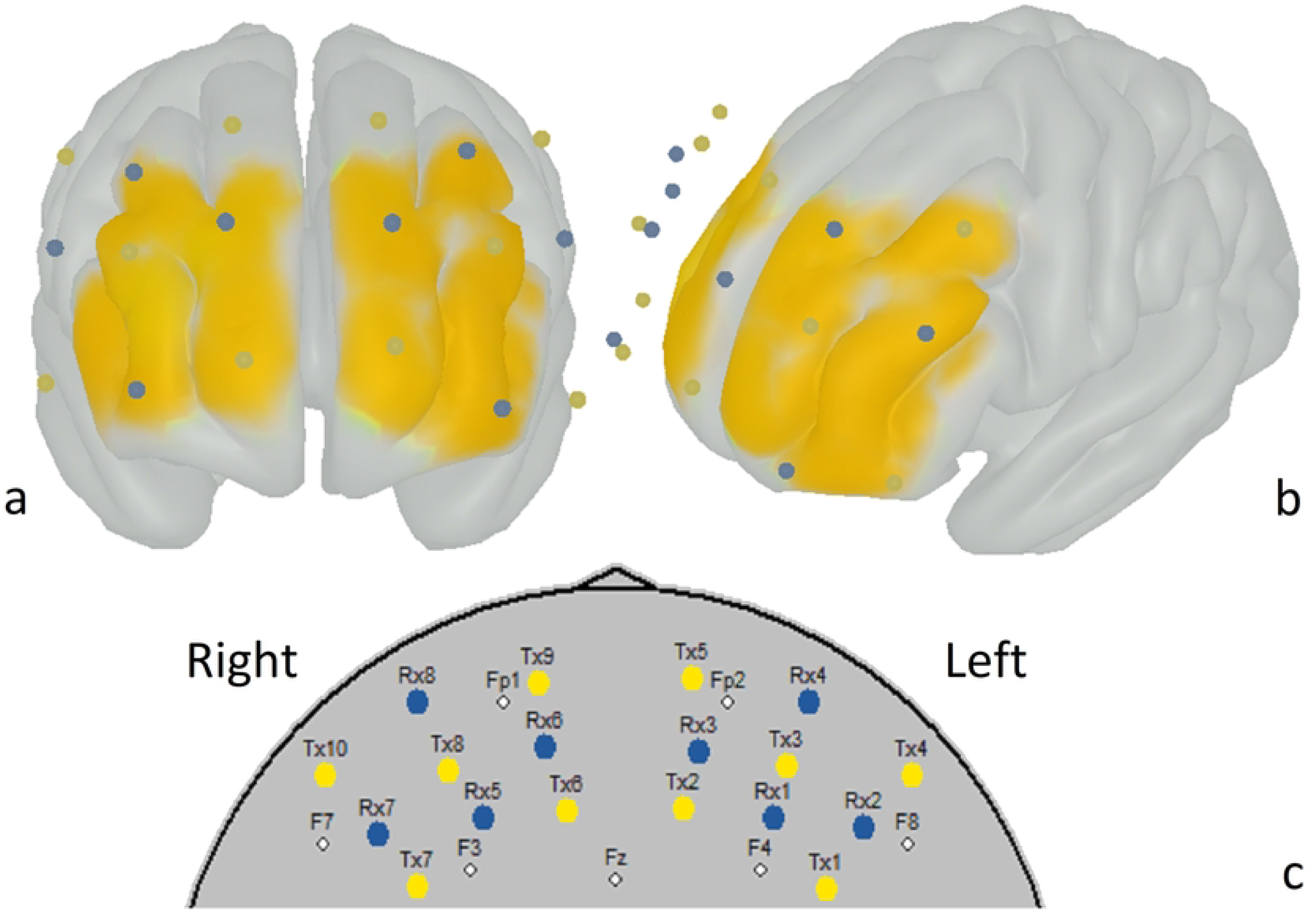
fNIRS optodes projection onto the frontal area. Blue dots represent transmitters, yellow dots represent receivers a) Anterior view b) Left front dorsal view c) 2D topographic view with location of transmitters and receivers regarding the international 10-20 system.

### Signals preprocessing

#### Electrodermal activity

To preprocess the EDA signals, we exported the E4 data into CSV files. The raw signal contains missing data due to sensor detachment which occurred randomly throughout the measurement sessions. In some cases, the detachment last for a few seconds, making it impossible to denoise solely with simple filtering techniques. Correspondently, we proposed an original measurement error detection algorithm. The proposed algorithm detects measurement errors by identifying steep changes in the EDA signals, such as those greater than 0.01 μS, which are outside the allowed range. The algorithm then examines the record until the measured EDA returns to the permitted range. When an error is observed in consequential timestamps, the allowed range is linearly increased by a factor equal to the difference between observations in a valid sequence. After applying this original algorithm, we interpolated the missing data using linear interpolation. For further removal of the high-frequency noise and smoothen the EDA signal, we applied a low-pass Butterworth filter with cut off frequency of 0.5 Hz [26]. Subsequently, all participants’ data were normalized using min-max scaling to ensure that the values fell within the 0-1 range. In the next step, the EDA values were averaged for each stage. Time stamps for EDA and fNIRS measurements were synchronized for each stage.

#### Cortical hemodynamics

The fNIRS signals were exported from the original OxySoft software, version 3, in the EDF format. The relative concentration changes in oxygenated hemoglobin (ΔOHb) and deoxygenated hemoglobin (ΔHHb) were computed using the Beer-Lambert law with an age-adjusted differential path length.

In the next step, we removed channels with poor signal quality using the Scalp Coupling Index (SCI) method [27]. We started by applying a band-pass filter (0.7–1.5 Hz) to each channel at both wavelengths to keep only the heartbeat components. After filtering, we normalized the signals to even out their amplitudes. We then calculated the zero-lag cross-correlation between the signals in each channel, which we called the SCI, to measure the signal-to-noise ratio. Channels with an SCI value below 0.75 were considered poor and were removed from further analysis. The good quality signals were band-pass filtered between 0.02 and 0.18 Hz to remove high-frequency components including cardiac and respiratory noises. The ΔOHb and ΔHHb values were averaged for each stage. The following stages were indicated: OS “baseline” (5 min); SA “anticipation” (5 min); AB “stress-A” (3 min); BC “stress-B” (4 min); CD “stress-C” (3 min); DE “recovery” (5 min).

### Statistical analysis

We performed paired t-tests to examine whether there were significant differences in the mean salivary cortisol before and after the IMPRO protocol. We also conducted a series of paired t-tests on the EDA signals using stage-by-stage comparison. To evaluate channel-wise hemodynamic changes in PFC we used a paired t-test to compare level of ΔOHb and ΔHHb between baseline and average stress. Next, we compared baseline with three IMPRO tasks “stress-A”, “stress-B”, “stress-C” and recovery stage correspondently. As suggested by previous studies [28, 29], we used the t-values for visualization of changes for each stage.

In addition, we investigated frontal asymmetry as a possible indicator of stress response. For this purpose, we performed two-way repeated measurement ANOVA for right-left groups of channels. Each participant’s fNIRS data were grouped and averaged for the right (channels 7, 9, 10, 12) and left (channels 14, 16, 17, 19) hemispheres (Fig 4). ANOVA tests were performed independently for oxy- and deoxyhemoglobin values. We verified the main effects of the within-group factor (stage) and the between-group factor (hemisphere).

**Fig 4.**
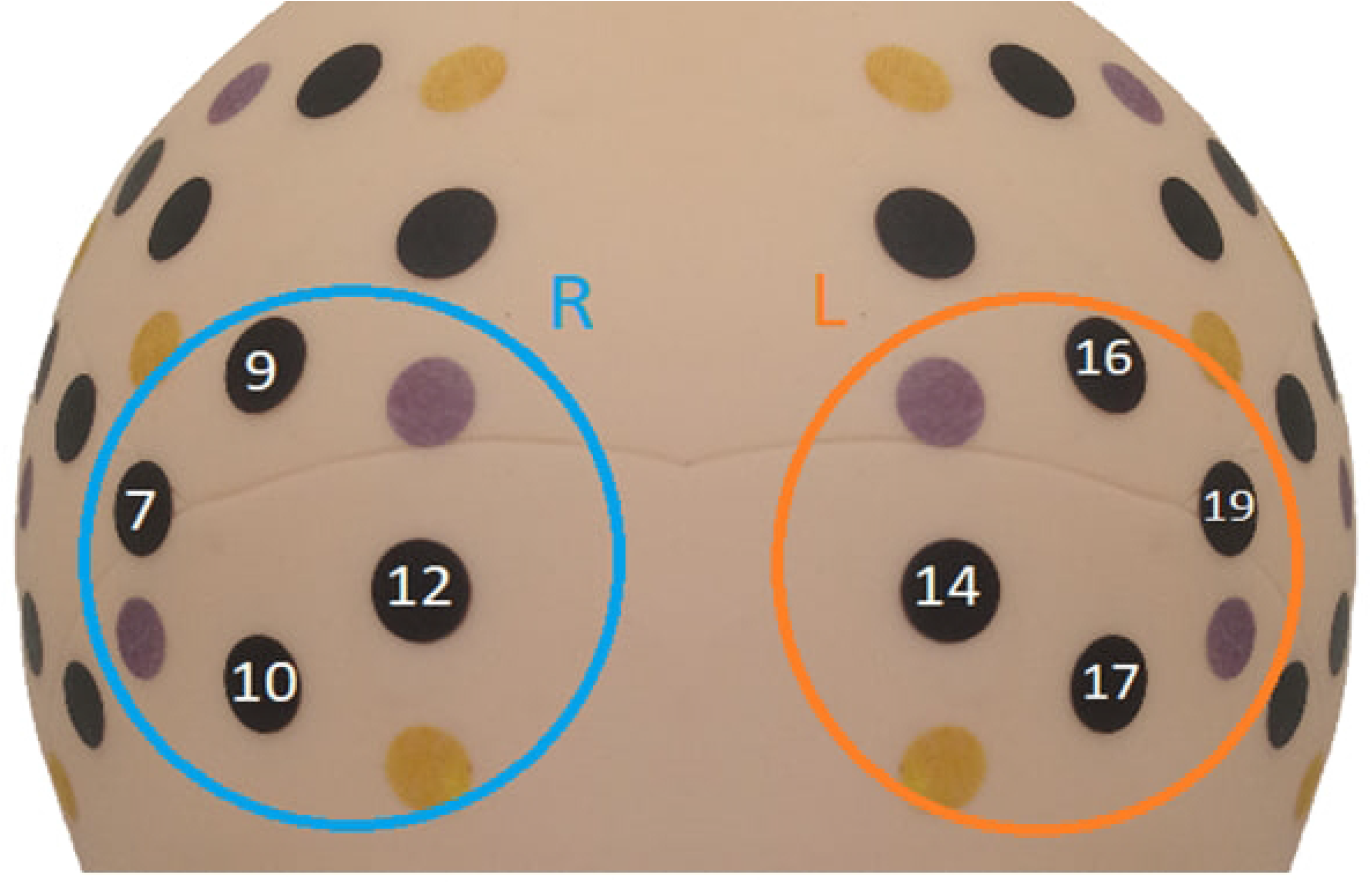
Grouping of the fNIRS channels regarding the frontal asymmetry investigation.

Finally, we compared mean values for each sub-stage and visualized significant changes and date-trends for EDA and fNIRS values observed throughout the experiment, including rest (baseline), anticipation, IMPRO-stress (free improvisation, random words challenge, arithmetic load), and recovery stages.

## Results

As shown in Fig. 5, the t-test showed a significant increase in the salivary cortisol levels in response to the IMPRO stress tasks (t = −3.4, p = 0.0013). The cortisol levels mean were 2.68 (SD=0.99) before and 3.54 (SD=1.25) after the IMPRO, respectively.

**Fig 5.**
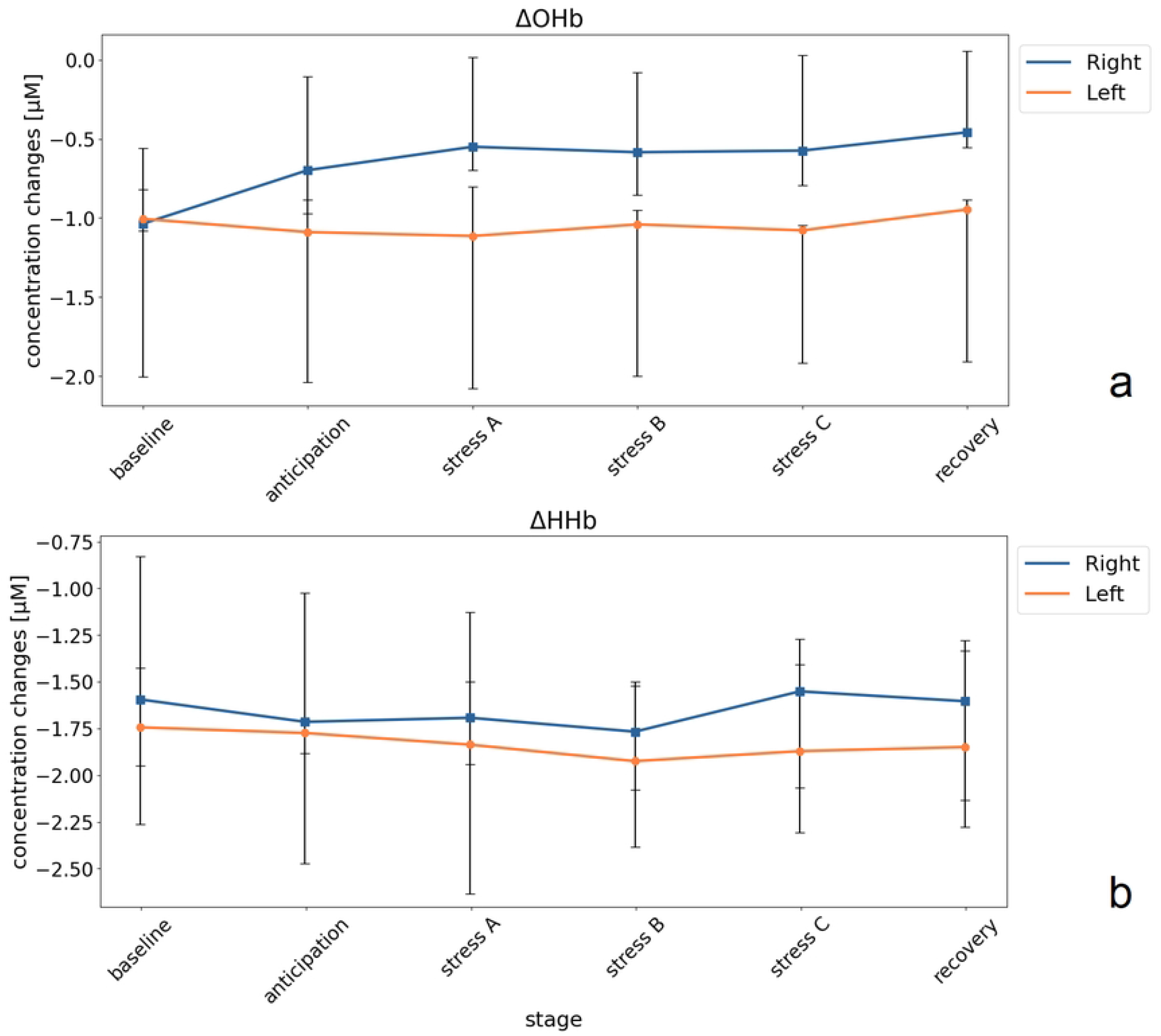
Salivary cortisol levels at the baseline and in response to the IMPRO stress tasks.

Results of the pairwise between-phase t-test on the EDA data are shown in Fig. 6. There were no significant differences between the baseline and anticipation stages. However, there was a significant increase in the EDA levels (t=-4.9; p=0.0003) in response to the sub-stage “stress A”. Skin conductance further increased during sub-stage “stress B”, random words improvisation (t=-2.8; p=0.019). No significant differences were found between sub-stages “stress B” and “stress C”, revealing no further increase in EDA during improvisation with arithmetic load. Then, a significant decrease was found during the recovery phase compared to the “stress C” stage (t=4.2; p=0.001).

**Fig 6.**
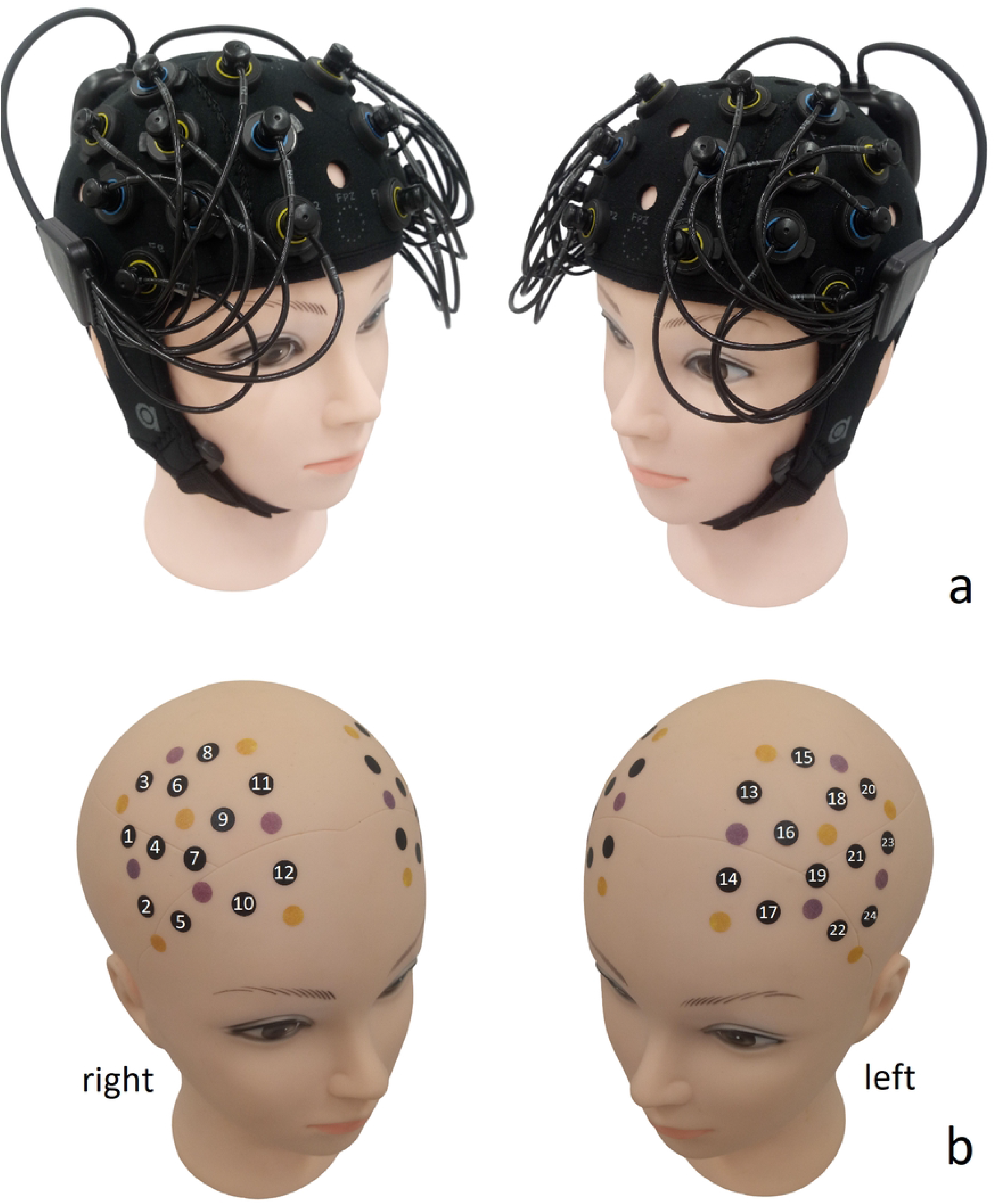
Electrodermal activity. Signal were normalized between 0-1, and averaged across six sub-stages for all subjects (**p<0,01, *p<0,05)

As to cortical hemodynamics, we first examined the difference between average baseline and average stress for all channels. ΔOHb signals increased during improvisation in the three channels: channel 10 (t=-2.2; p=0.0421), channel 12 (t=-2.2; p= 0.0169), and channel 17 (t= −2.4, p = 0.031), ΔHHb signals showed a significant decrease in channel 22 (t-value = 2.5, p = 0.024) (Table 2).

**Table 2.** Channels with significant difference between baseline and average stress stages, showing t-test channel-wise results.

| Oxy/Deoxy hemoglobin | Channel | t-value | p-value | Brodmann area | Hemisphere |
| --- | --- | --- | --- | --- | --- |
| $\Delta\text{OHb}$ | 10 | -2.23 | 0.0421 | 10 | Right |
|  | 12 | -2.71 | 0.0169 | 10 | Right |
|  | 17 | -2.41 | 0.0310 | 10 | Left |
| $\Delta\text{HHb}$ | 22 | 2.55 | 0.0240 | 45 | Left |

Comparison across substages showed significant differences in the same channels (10,12,17) with the most pronounced increase in oxyhemoglobin levels in the initial phase of improvisation (stress A), as shown in Fig 7.

**Fig 7.**
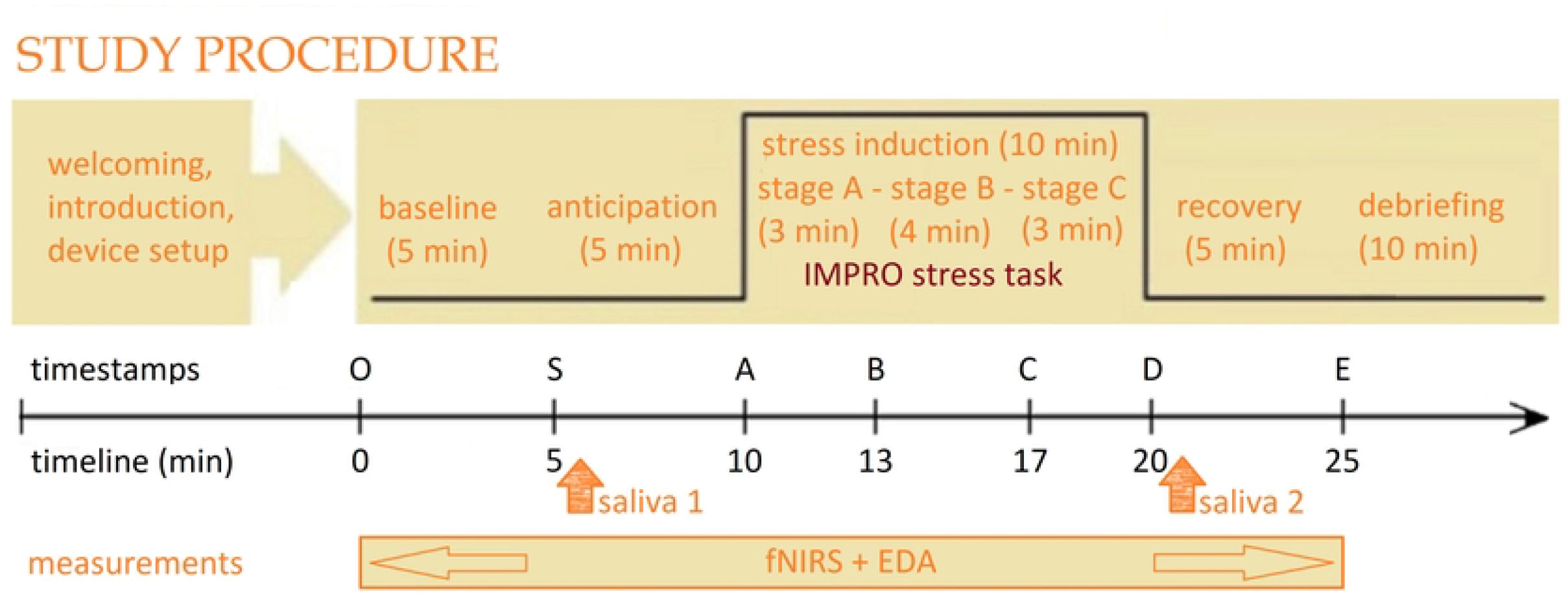
fNIRS optodes projected onto the frontal area.

The ANOVA test for ΔOHb showed significant changes in oxyhemoglobin concentration in response to the IMPRO tasks (p<0.05). It was also found that the oxyhemoglobin level growth was more pronounced in the right hemisphere (Table 3). The variations in mean fNIRS levels for each sub-stage for the right and left group of channels are shown in Fig 8.

**Table 3.** Stage by Hemisphere two-way repeated measurement ANOVA for mean levels of oxygenated (ΔOHb) and deoxygenated (ΔHHb) hemoglobin.

| Oxy/Deoxy hemoglobin | Factor | F-value | p-value uncorrected | p-value corrected |
| --- | --- | --- | --- | --- |
| OHb | hemisphere | 7.77 | 0.0191 | 0.0191* |
|  | stage | 2.41 | 0.0486 | 0.0987* |
|  | hemisphere * stage | 3.84 | 0.005 | 0.0435* |
| HHb | hemisphere | 2.70 | 0.1238 | 0.1238 |
|  | stage | 0.73 | 0.5984 | 0.5166 |
|  | hemisphere * stage | 0.06 | 0.9969 | 0.9453 |

**Fig 8.**
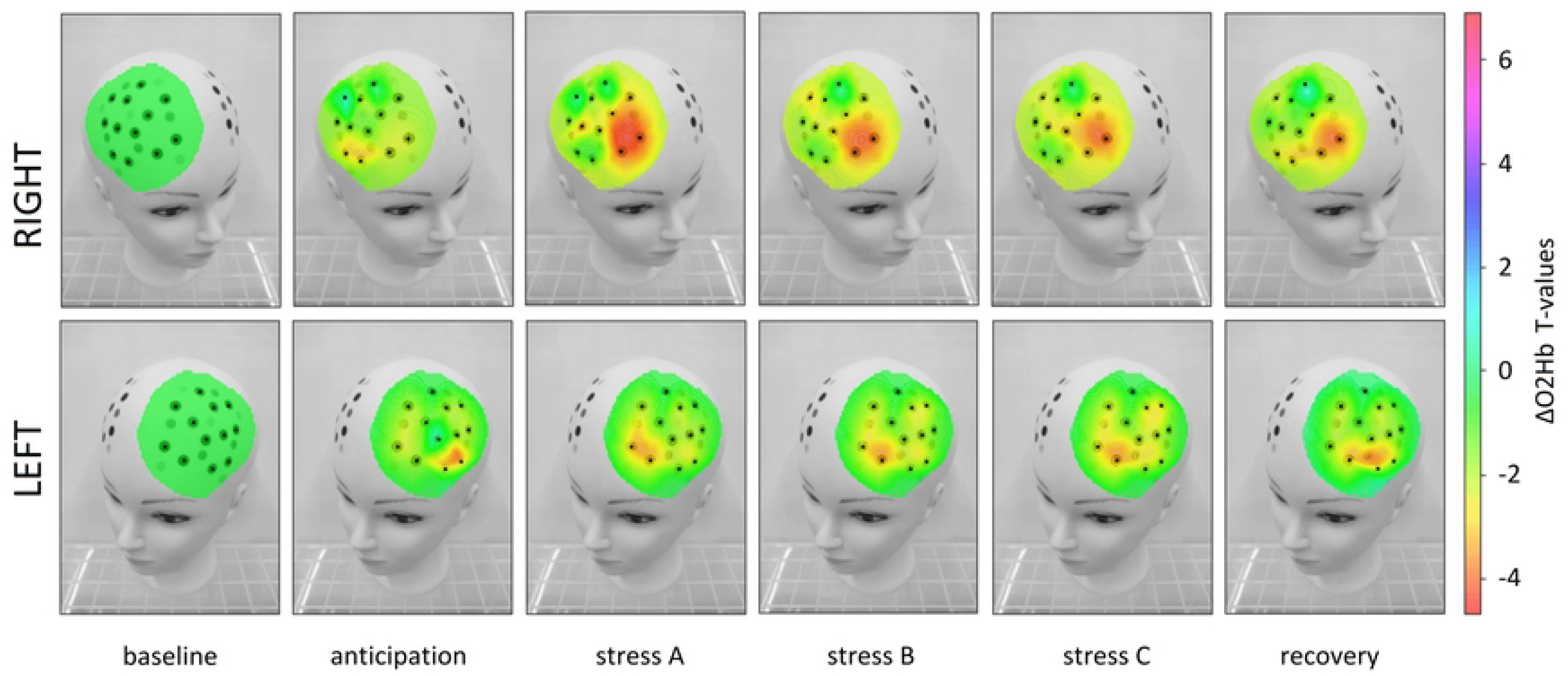
Hemoglobin concentration changes by stage and hemisphere. a) mean values for oxygenated blood (ΔOHb) b) mean values for deoxygenated blood (ΔHHb)

## Discussion

### Validity of IMPRO stress induction protocol

The current study aimed to validate a novel improvisation-based stress induction protocol. The principal finding for the IMPRO protocol was a significant increase in cortisol levels following the stress induction. Cortisol, a hormone released by the adrenal glands in response to stress, plays a pivotal role in mobilizing the body’s resources to cope with perceived threats. The heightened cortisol levels indicate activation of the hypothalamic-pituitary-adrenal (HPA) axis, mirroring typical stress reactions observed in classical protocols like the Trier Social Stress Test [30, 31]. Furthermore, EDA served as a valuable indicator of autonomic arousal. The coherence between EDA findings and elevated cortisol levels provides strong support for the validity of both physiological markers in capturing the comprehensive stress response. Together, cortisol and EDA provide complementary insights into different aspects of how speech improvisation can affect performers’ bodies. Cortisol indicates the endocrine response to stress, affecting metabolism, immune function, and other bodily processes, while EDA reflects immediate sympathetic nervous system changes related to emotional arousal and attentional focus [32]. This finding underscores the validity of the proposed IMPRO protocol for stress induction.

In addition to its validity, our IMPRO protocol has several advantages compared to existing stress induction protocols. In terms of in-lab implementation, the IMPRO protocol is less resource-intensive than the TSST. It requires minimal preparation and fewer human resources, making it more accessible for a variety of research settings. This ease of use is particularly beneficial for studies with limited resources or those conducted outside of a laboratory environment. The IMPRO protocol boasts high ecological validity, closely mimicking real-life scenarios that require spontaneous thinking and adaptability. This realistic approach is more reflective of everyday stressors, making the findings more applicable to situations in the field. Additionally, the flexibility of the improvisational tasks allows for customization to specific populations and research needs, enhancing the personal relevance and impact of the stressor. Of particular importance in our protocol is the gamification element. Reducing the significance of the stressor stimulus by presenting the task in a game context. So, we provoke what is called *eustress*. In our experiment, the physiological response is maintained, while the potential negative impact on emotional vulnerability of the participants is mitigated, making the protocol less invasive and ethically acceptable for a wide range of age and social groups.

### Cortical responses during IMPRO stress induction

Our findings of the cortical responses as manifested in fNIRS measurements align with previous brain imaging studies on mental stress. The frontal asymmetry has been previously identified in several studies using fMRI, EEG, and more recently, fNIRS [25,32-34]. For example, Beraha et al. demonstrated region-specific lateralization in emotion processing supported by fMRI. This study supports the valence asymmetry hypothesis indicating positive affective stimulus processing is lateralized towards the left hemisphere in the medial prefrontal cortex, while negative affective stimulus processing shows right-lateralization in the dorsolateral prefrontal cortex and left-lateralization in the amygdala and uncus [33].

However, extended studies on hemispheric asymmetry shows that no single model of frontal asymmetry is unequivocally supported for mood processing, as results are contradictory across studies, with critical methodological issues such as sample composition, mediating third variables, and the interplay between mood and cognition needing further exploration [34]. More recent fNIRS studies showed that variations of the PFC response may depend on multiple preliminary factors, such as preliminary anxiety [25].

Besides, resting brain activity, when lateralized to the left or right, significantly predicts both prefrontal cortical responses and explicit appraisal of emotional stimuli [35].

Nevertheless, a large number of neuroimaging studies point to frontal asymmetry as one of the significant markers of affective regulation. Our study contributes to this body of work by employing speech improvisation, which uniquely combines social stress, cognitive load, and spontaneous performance. Unlike previous studies, our use of fNIRS allows for continuous monitoring of prefrontal cortex activity in a more naturalistic setting, offering new insights into the temporal dynamics of stress responses and their recovery patterns.

Our device connected through Bluetooth provides maximum freedom and mobility to participants and creates an experimental environment similar to a natural communication setting. Increased activity in a set of channels, especially oxyhemoglobin changes in the frontopolar cortex (Brodmann’s 10th area), as well as lateralization of the signal with more significant activation of the right hemisphere, is often associated with mental stress.

Moreover, some studies have noted the importance of PFC in shaping spontaneous performance and response flexibility [36]. Acknowledging the prefrontal cortex’s multifunctionality, we emphasize our primary objective was to track the stress response. Integrating multiple evidence, we can suggest the involvement of the stress response system at all levels. The use of wearable devices provided a non-invasive, real-time measure of physiological activity, offering a dynamic view of how stress affects the brain and body.

### Limitations

Despite the promising results, this study has several limitations. First, the protocol involves a mixture of different types of interventions and cognitive processes, including social stress, cognitive load, speech production, and improvisation (bearing in mind spontaneous performance without prior preparation). This multifaceted approach makes it challenging to isolate the specific contributions of each factor to the overall stress response. Inherent variability in improvisational tasks may lead to inconsistencies in the stress response, implies the need of further research to standardize the protocol while preserving its benefits. Second, the limited time allocated for recovery stage restricts our conclusions regarding the dynamics in the post-intervention period. The presented study includes only two measurements of cortisol levels in each experiment session, which does not capture the recovery phase following the stress tasks. Nonetheless, recovery can be demonstrated by other physiological signals such as EDA. Personal chronotype also may act as a confounding factor. Cortisol changes related to the daily circadian rhythms may interfere with changes related to the task of interest. Third, the use of fNIRS limits our neuroimaging to superficial cortical structures, such as the prefrontal cortex, while deeper brain regions involved in stress responses remain inaccessible. Additionally, the study’s fNIRS setup utilized 24 channels focusing on the frontal area, restricting our ability to explore brain connectivity comprehensively. Lastly, future studies should investigate the psychological effects of the protocol, including subjective measures of stress and anxiety, to complement the physiological data.

### Further research

Further studies may specify those neural circuits that enable speech improvisation flow by extending the number of channels to cover additional cortical areas. Furthermore, although with necessary limitations in movement freedom, spontaneous speaking can be modified for fMRI studies for further examination of brain activity with capturing deeper brain structures. Secondly, this technique can be adapted for individualized stress-management practices outside the laboratory. We can conduct exposure-type therapeutic interventions to train the ability to act spontaneously under stressful conditions with high uncertainty levels while reducing anxiety through behavioral training. Moreover, integrating wearable device tracking into mental health practices could enhance the effectiveness of stress management, both for clinical treatment and daily well-being support.

## Conclusion

This study demonstrates that an improvisation-based stress induction protocol, validated through fNIRS neuroimaging, is an effective alternative to traditional methods like the TSST. Its high ecological validity, flexibility, and practical advantages make it a valuable tool for stress research and management. The potential integration of portable fNIRS and wearable devices heralds a new era in personalized stress management, offering promising solutions for improving mental health and well-being. Future research should continue to refine this protocol and explore its applications in diverse populations and a wide range of settings.

